# The pulvinar regulates plasticity in human visual cortex

**DOI:** 10.1101/2025.02.24.639829

**Authors:** Miriam Acquafredda, Jan W. Kurzawski, Laura Biagi, Michela Tosetti, Maria Concetta Morrone, Paola Binda

## Abstract

In normally sighted human adults, two-hours of monocular deprivation is sufficient to transiently alter ocular dominance. Here we show that this is associated with a reduction of functional connectivity between the pulvinar and early visual cortex, selective for the pulvinar-to-V1 directionality. Across participants, the strength of the pulvinar-to-V1 connectivity was negatively correlated with the ocular dominance shift, implying less plasticity in participants with stronger influence of the pulvinar over V1. Our results support a revised model of adult V1 plasticity, where short-term reorganization is gated by modulatory signals relayed by the pulvinar.

**Teaser:** Short-term monocular deprivation triggers a reorganization of the visual processing network, measured with 7TfMRI in human adults.

## Introduction

The potential for long-term plastic change in the sensory brain declines with age, probably through active stabilization of the sensory circuitry (*1*). Nevertheless, adult sensory systems retain the potential for short-term plasticity (*2, 3*): blocking vision in one eye for a few hours in human adults induces a transient boost of the deprived eye (*4, 5*). This response modulation was observed using fMRI in primary visual cortex V1 (*6*), but not in the Lateral Geniculate Nucleus (*7*), indicating that the modulation of cortical activity was not inherited from earlier stages. Visual responses were also modulated in an adjacent thalamic nucleus, the pulvinar (*7*); whether this pulvinar modulation was a product of the V1 signal-change or involved in its generation remained an open question. The homeostatic response reflects a gain-change of the deprived eye representation, which could serve to counterbalance the reduced stimulation (*8*). However, there is recent evidence that the same boost can be achieved without blocking stimulation of either eye, merely manipulating attention and multimodal context (*9–12*). This suggests a radically novel interpretation of adult plasticity, where top-down multimodal signals play a key role.

One main pathway through which multimodal signals reach V1 is via the pulvinar, which modulates activity through recurrent cortico-thalamo-cortical loops (*13, 14*). These loops connect distant sensory, motor and cognitive areas and may underly predictive coding (*15–17*): by collecting multimodal signals, the pulvinar may carry predictions about the upcoming sensory events to low-level sensory areas, including V1 (*18, 19*). In line with this model, there is evidence that reducing the pulvinar influence over V1 permits the updating of perceptual representations, i.e. perceptual learning (*20*).

Based on this evidence, we hypothesize that the pulvinar exerts a stabilizing influence over the primary sensory cortex, and that reducing this influence may open an opportunity for plasticity – even for a very stable property like ocular dominance. To test this idea, we used high resolution 7T-fMRI and asked whether short-term monocular deprivation impacts the pulvinar-V1 connectivity.

## Results

### Short-term monocular deprivation alters pulvino – cortical functional connectivity

We first confirmed the deprivation effect in 22 normally sighted adults. After the two-hour monocular deprivation, ocular dominance, measured with a short binocular rivalry test immediately before MRI scanning, shifted in favor of the deprived eye by 10% on average (from 55% to 65% dominance, Figure 3A).

Next we measured the resting-state (eyes closed) functional connectivity between the pulvinar (atlas-based ROI definition (*21*)) and the cortex; this was strong in all sensory regions, including occipital visual areas, temporal auditory areas and central somatosensory areas (Figure 1A, left) in four highly significant clusters (continuous white lines), consistent with the multimodality of the pulvinar (*22, 23*). After monocular deprivation (Figure 1A, middle), there was a dramatic reduction of pulvinar connectivity with the occipital cortex, including the primary visual cortex V1 and surrounding early visual areas (Figure 1A, right, displaying the post-pre deprivation difference) as well as fewer and sparser reductions in the temporal and parietal cortex.

**Figure 1.**
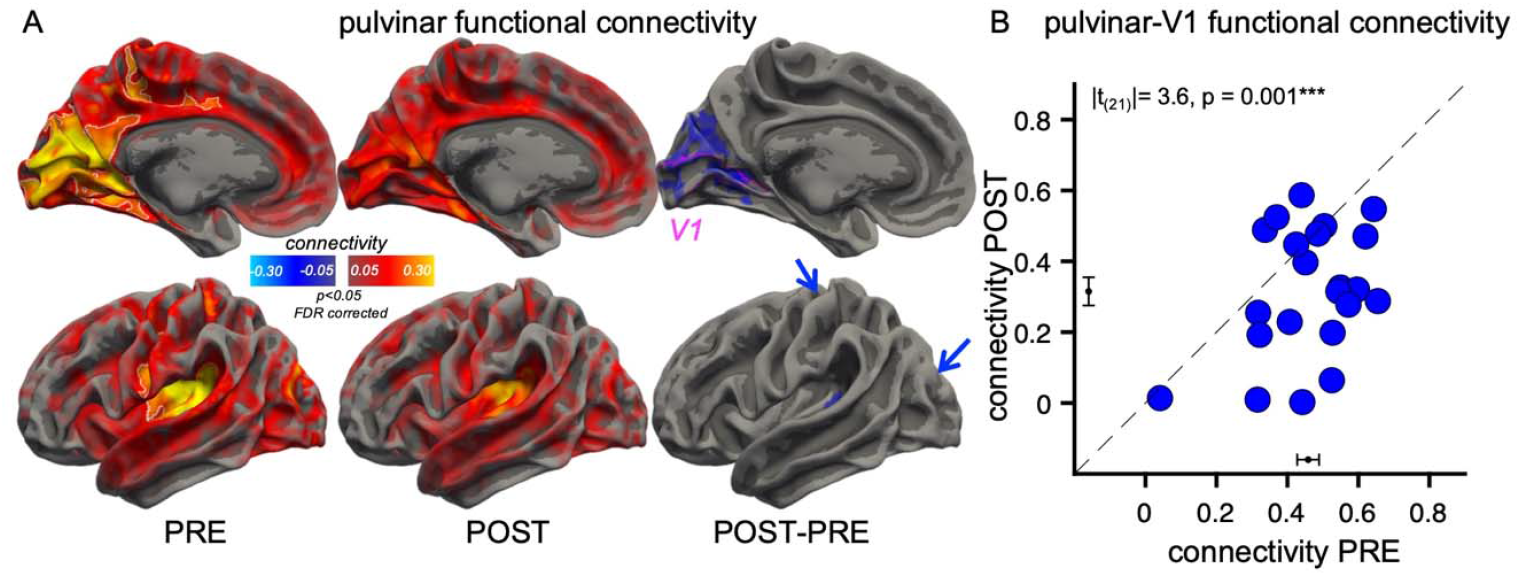
Functional connectivity of the pulvinar, measured with resting-state 7T-fMRI. A: Functional connectivity of the pulvinar, estimated from resting-state fMRI (TR = 3s, full brain coverage) and mapped over the cortical surface (medial and lateral view; maps are averaged across hemispheres and projected on a symmetrical template of the cortical surface). The leftmost and central maps show the connectivity before and after two hours of monocular deprivation; the rightmost maps show the connectivity change, showing significant reductions in occipital visual areas (mainly mesial and one postero-lateral, not visible and signaled by the posterior blue arrow); there was also a connectivity reduction in two small areas of the superior temporal gyrus (visible in the lateral view) and the central sulcus (signaled by the anterior blue arrow). Connectivity was estimated as the Pearson’s r coefficient between the fMRI timeseries in the pulvinar and the time-series of each cortical voxel, computed after nuisance regression and concatenation across N=22 participants. Maps are thresholded at |r| > 0.05 and p < 0.05 FDR corrected, and cluster corrected (50 vertices). The white outlines in the pre-deprivation map identify the clusters of regions where connectivity is highly reliable (|r| > 0.20 and 500 vertices); the pink outline in the map of connectivity differences shows the V1 ROI. B: Functional connectivity between the pulvinar and V1 (atlas-based definition, see pink outline in Panel A) before and after deprivation, for the individual participants. Pearson’s r values were Fisher transformed to allow for statistical comparisons; the black error bars near the axes show the mean +/− S.E.M. connectivity: 0.46 ± 0.03 before deprivation reduces to 0.32 ± 0.04 after deprivation. The text inset reports the results of the paired t-test comparing functional connectivity before and after deprivation (Cohen’s d = 0.80).

This connectivity reduction after deprivation was confirmed by calculating functional connectivity for the individual participants (Figure 1B), from the average time series of two a-priori selected Regions of Interest: pulvinar (*21*) (illustrated in Figure 2B) and V1(*24*) (illustrated by the pink outline in Figure 1A and the pink region in Figure 2A). The connectivity change was not accompanied by changes in BOLD modulations, as the overall power of the Fourier spectrum of the signals in each region of interest remained comparable before and after deprivation in all regions of interest (V1: |t_(21)_|= 0.3, p = 0.803, Cohen’s d = 0.05; pulvinar: |t_(21)_|= 1.4, p = 0.173, Cohen’s d = 0.30). The change in pulvinar connectivity extended beyond V1 to several other visual areas (Supplementary Figure S3A), and to few areas of the superior temporal gyrus and central sulcus (Supplementary Figure S3A). In contrast with the dramatic reduction of pulvinar connectivity, no change was observed for LGN connectivity (*25*) (Supplementary Figure S1A-B), nor for connectivity between V1 and the other occipital areas (Supplementary Figure S2). There was, however, a sparse reduction of V1 connectivity with high-level areas, distributed across the superior temporal gyrus, the intraparietal sulcus, the premotor cortex and the cingulate gyrus (Supplementary Figure S2).

**Figure 2.**
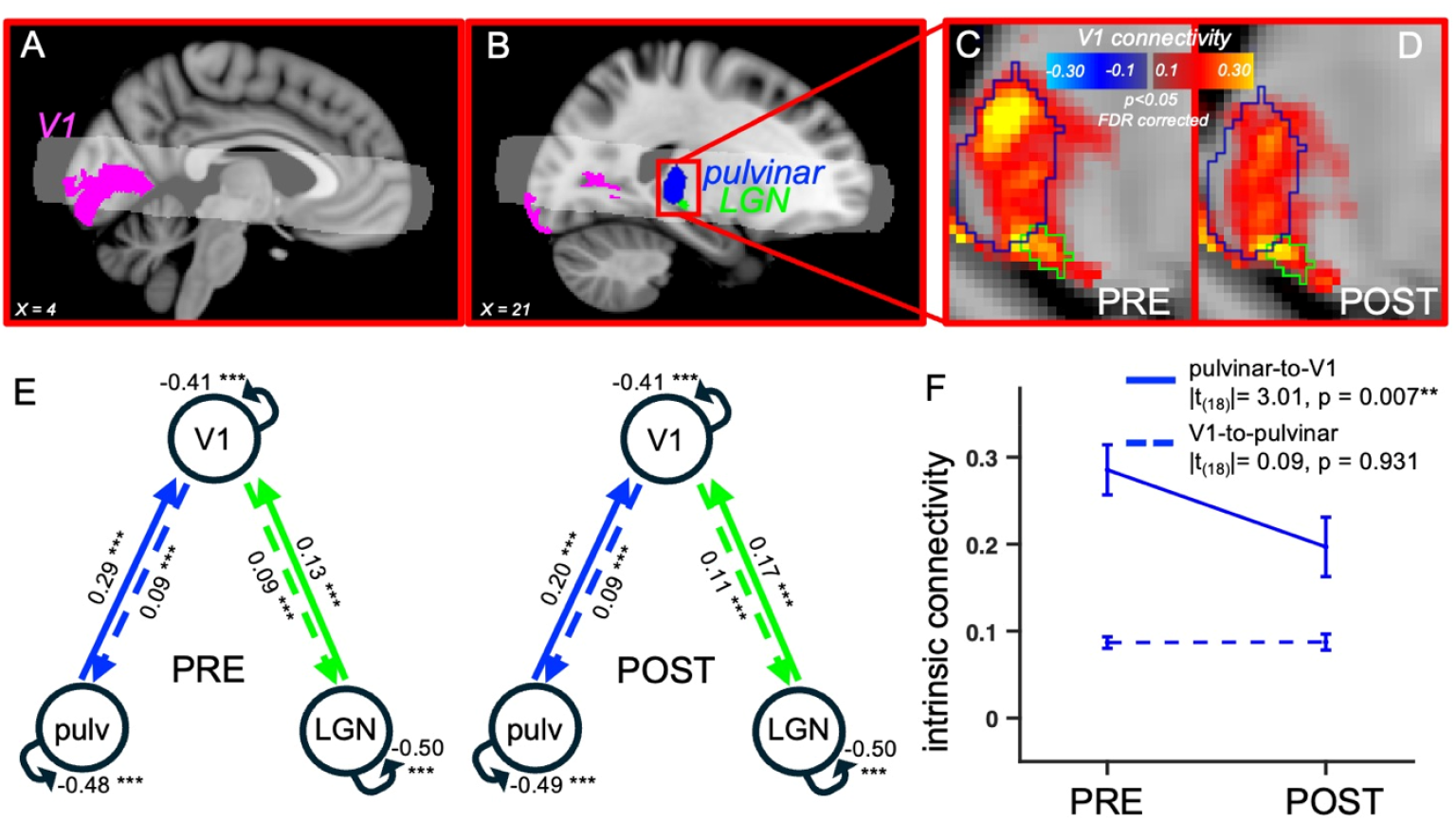
Thalamus-V1 functional connectivity and effective connectivity measured with Dynamic Causal Modelling of the resting-state 7T-fMRI time-series. A-B: Partial volume covered by a high-temporal resolution fMRI (TR = 1s, partial volume: white shading shows spatial coverage across 50% of our acquisitions) used for measuring connectivity between V1 (defined as the intersection between the atlas-based V1 region of interest (24), in pink, and the coverage afforded by our sequence) and the thalamic regions pulvinar and LGN (atlas-based definitions (21, 25), blue and green masks), visualized in two different sagittal slices (x-MNI coordinates shown as text insets). C-D: Functional connectivity between V1 and the thalamus, before and after monocular deprivation. Connectivity was estimated as the Pearson’s r coefficient between the fMRI time-series in V1 and the timeseries of each voxel in the thalamus, computed after nuisance regression and concatenation across N=19 participants; maps are thresholded at |r| > 0.05 and p < 0.05 FDR corrected. See Supplementary Figure S5A-B for a coronal view of the V1 connectivity profile within the pulvinar ROI. E: Average effective connectivity across subjects in the network including bidirectional connections between V1 and the two thalamic regions: pulvinar and LGN; Numbers show intrinsic connectivity (A-matrix in rDCM) and associated statistical significance (after FDR correction: *** for p < 0.001, ** for p < 0.01, * for p < 0.05, ns for p >= 0.05), independently estimated before and after monocular deprivation. F: Change of effective connectivity between the pulvinar and V1, shown as mean (same numbers as in E) and S.E.M across participants. Text insets report the post-hoc t-tests comparing effective connectivity values before and after deprivation for the two directionalities (after FDR correction: **p < 0.01, * p < 0.05; Supplementary Figure S4A shows results for an alternative DCM network, assuming reciprocal connectivity between all areas (including LGN and pulvinar) and characterized by a worse fit to the data.

### The functional connectivity reduction is specific for the pulvinar-to-V1 directionality

To estimate the directionality of the pulvinar-V1 connectivity change (feedforward vs. feedback), we acquired a second resting-state fMRI dataset, with higher temporal resolution (TR = 1 second, covering V1 and the thalamus, Figure 2A-B). The V1 functional connectivity with the thalamus (Figure 2C) shows strong connectivity both in the atlas-based definition of LGN (*25*), and in the atlas-based definition of the pulvinar (*21*). Comparison of the V1 connectivity map before and after deprivation (Figure 2C-D) suggests a selective reduction in the pulvinar, but not LGN. We applied Dynamic Causal Modelling (*26*) to estimate bidirectional effective connectivity among the three regions: V1, LGN and pulvinar. We tested two models, one fully connected (Supplementary Figure S4A), and a more anatomically plausible model with no direct connections between LGN and Pulvinar, shown in Figure 2E. The latter showed a better fit to the data than the fully connected model (negative free energy, |t_(18)_| = 88.7, p < 0.001, Cohen’s d = 20.3). Effective connectivity estimates from this model on the individual participants’ data (Figure 2E-F) shows that monocular deprivation selectively reduced the pulvinar-to-V1 connectivity leaving the LGN-to-V1 connectivity and the connectivity from V1 to either of the thalamic regions unaffected. A 2×2 ANOVA with factors time (pre vs. post deprivation) and directionality (thalamus-to-V1 vs. V1-to-thalamus) showed a significant interaction for the effective connectivity between the pulvinar and V1 (F_(1,18)_ = 8.30, p = 0.010, η^2^_partial_ = 0.32), not between LGN and V1 (F_(1,18)_ = 0.2, p = 0.638, η^2^_partial_ = 0.01); post-hoc t-tests comparing effective connectivity values before and after deprivation for the two directionalities showed significant reduction of pulvinar-to-V1 connectivity (|t_(18)_|= 3.01, p = 0.007, Cohen’s d = 0.69), no change of V1-to-pulvinar connectivity (|t_(18)_|= 0.09, p = 0.931, Cohen’s d = 0.02) and no change for LGN-toV1 (|t_(18)_|= 0.75, p = 0.464, Cohen’s d = 0.17) or V1-to-LGN connectivity (|t_(18)_|= 0.51, p = 0.617, Cohen’s d = 0.12). Similar results hold for the fully connected model (Supplementary Figure S4A), although this model assigns significant connectivity values to the LGN-pulvinar connection before deprivation, which is not supported by anatomical evidence (e.g.*27*). The DCM-based conclusions are also supported by a more direct cross-correlation analysis of the BOLD timeseries (shown in Supplementary Figure S4B).

The DCM estimate of pulvinar-to-V1 connectivity was related to ocular dominance measured behaviorally. The variability across-participants in the strength of the pulvinar-to-V1 connectivity before deprivation reliably predicted the change of ocular dominance with monocular deprivation (Figure 3B). The correlation is negative: subjects with stronger pulvinar influence over V1 showed less ocular dominance plasticity. We did not observe a reliable correlation between connectivity changes and the change of ocular dominance (r_(18)_ = −0.05, p = 0.834), probably reflecting the increase in error associated with the difference between the pre- and post-deprivation noisy estimates of connectivity.

**Figure 3.**
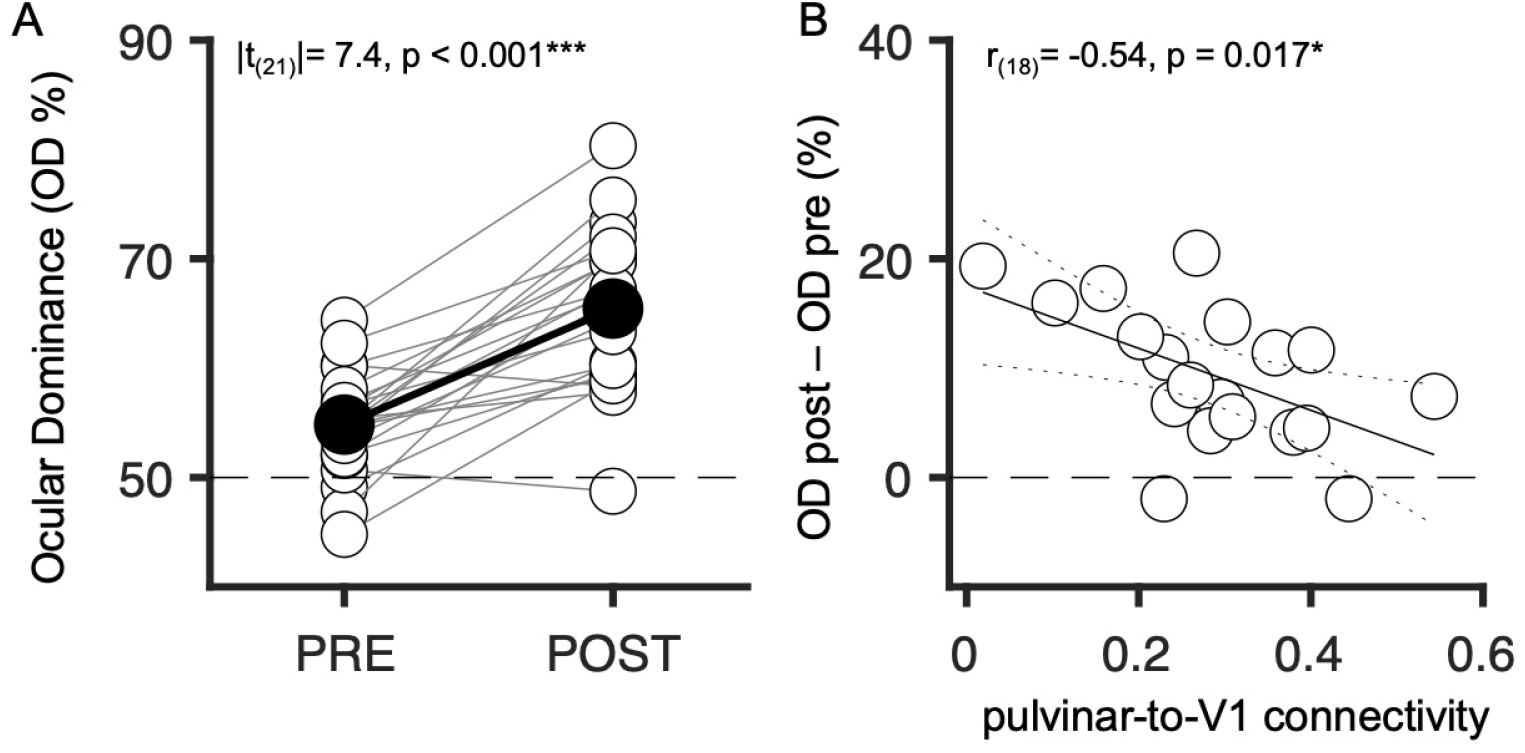
Ocular dominance plasticity correlates with pulvinar-to-V1 connectivity. A: Ocular dominance measured with binocular rivalry before and after the two-hour monocular deprivation, shifting in favor of the deprived eye. Each empty symbol is one participant; pre- and post-deprivation values are connected by thin lines. Filled symbols and the thick line show the average across participants and the paired comparison across time-points is reported in the text inset. B: In the subset of 19 participants included in the DCM analysis, the ocular dominance shift was negatively correlated with the pulvinar-to-V1 connectivity estimated with DCM from pre-deprivation acquisitions. The text inset gives the Pearson’s correlation coefficient and associated p-value. Continuous and dotted lines show the best-fitting linear function and the 95% confidence bands.

The V1 connectivity map in the pulvinar and its change with deprivation showed at least two distinguishable foci (Supplementary Figure S5A-B). To gain further insight into possible subdivision of pulvinar region (Figure 2C-D), we recorded visual BOLD responses in two conditions differing in voluntary attention allocation (Supplementary Figure S5C). We used the same stimuli that reliably mapped thalamic activations (*7*); these were presented either in passive viewing or while the subject allocated attention to the stimulus area to detect small peripheral color changes. As expected, passive viewing elicited a cluster of activity within the ventral portion of the pulvinar, and attending to the stimuli elicited additional activity in a more dorsal portion. We used these activity maps to define a ventral visual pulvinar (cyan outline in Supplementary Figure S5A-C) and a relatively dorsal pulvinar modulated by attention allocation (magenta outline). The union of these two sub-ROIs covered about half the anatomically defined pulvinar *(21)*, and the connectivity of this region with V1 was reliably stronger than in the remaining half of the pulvinar (|t_(18)_|= 5.7, p < 0.001, Cohen’s d = 1.30). By applying our DCM-based approach to each sub-region, we observed the reduction of pulvinar-to-V1 connectivity was present only for the relatively dorsal region (Supplementary Figure S5D-E), supporting the implication of a high-level mechanism.

### Short-term monocular deprivation does not impact thalamo-cortical structural connectivity

We also assessed whether these functional connectivity changes are associated with changes in structural connectivity. We acquired Diffusion-Weighted MR Images during the same sessions as the resting-state scans. Figure 4 shows that thalamus-V1 structural connectivity was unaffected by monocular deprivation.

**Figure 4.**
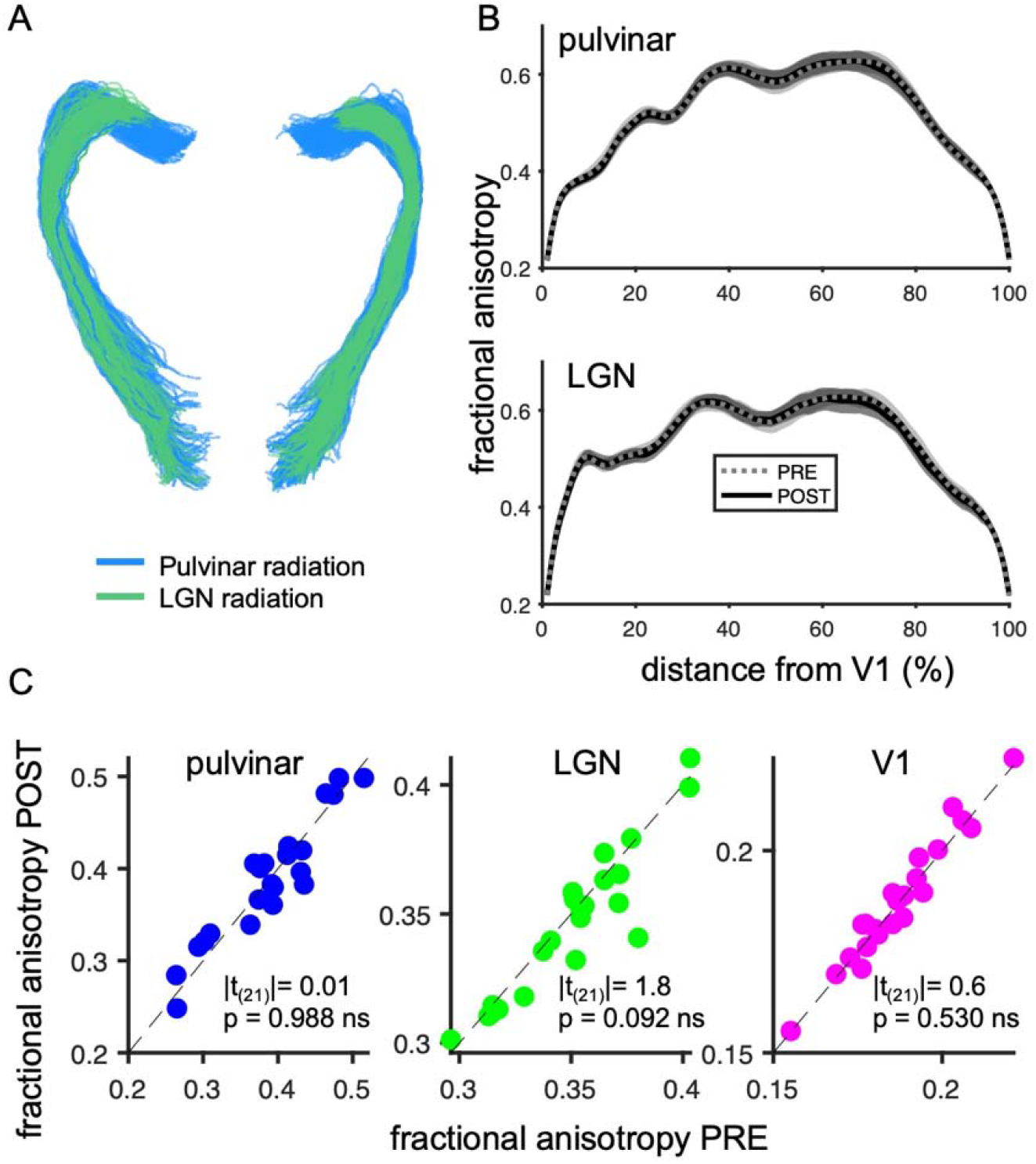
Structural connectivity between the thalamus and V1, measured with Diffusion Weighted Imaging. A: Fiber-tracking (visualized in DSI-STUDIO) for a representative participant, mapping the structural connectivity between V1 and two thalamic regions: LGN and pulvinar. B: Fractional anisotropy values as a function of nodes along the tracts (averaged across hemispheres), measured before (dashed gray lines) and after monocular deprivation (continuous black lines); shaded areas show 1 S.E.M. across participants. No difference was detected before vs. after monocular deprivation (paired t-tests, all p > 0.05 FDR corrected). C: Fractional anisotropy before vs. after monocular deprivation within each region, for the individual participants. Text insets report the results of paired t-tests comparing fractional anisotropy before and after deprivation (Cohen’s d = 0.00, 0.38 and 0.14 for pulvinar, LGN and V1).

## Discussion

Our results show that two hours of monocular deprivation reshapes the reciprocal interactions between visual brain regions, decreasing the influence of the pulvinar on resting-state V1 activity. In contrast, monocular deprivation did not reduce the LGN bottom-up sensory drive to V1.

Functional connectivity between the pulvinar and the cortex was reduced in both V1 and other visual areas, mainly in the dorsal mesial occipital cortex, with sparser connectivity changes in the auditory and somatosensory cortex. In V1, where effective connectivity could be estimated at high temporal resolution, the reduction was selective for the pulvinar-to-cortex connectivity, while connectivity in the cortex-to-pulvinar direction was remarkably stable. A similar trend was observed for the only other cortical area covered by our high-temporal resolution acquisitions that showed significantly reduced functional connectivity with the pulvinar, corresponding with the auditory Belt area. We also analyzed the topography of functional connectivity between V1 and individual voxels within the pulvinar and found that connectivity changes were primarily localized within a dorsal sub-region of the pulvinar. This region was defined by its modulation with attention, implicating the fronto-parietal network (*13, 28*) and top-down signals in the monocular deprivation effect. The topography of the effects, both across the cortex and within the pulvinar, are consistent with modulation by top-down contextual signals accompanying plasticity in early visual cortex.

The short-term effects of monocular deprivation in adults that we study here are very different from the long-term effects observed during development, when permanent suppression of the deprived-eye representation is accompanied by reduced bottom-up drive from LGN (*1*). The counterintuitive and opposite effect elicited by short-term deprivation (deprived-eye enhancement) is often interpreted as a homeostatic response to the reduced sensory input in the deprived eye (*8, 29*). Our results add to previous evidence (*7*) indicating that the gain-change is not accompanied by a modulation of the bottom-up input from LGN; they suggest that it is associated with decreased top-down influences conveyed through the pulvinar – primarily a dorsal sub-region of the pulvinar. We find that the strength of the pulvinar-to-V1 connectivity predicts the participants’ behavioral susceptibility to plastic changes, demonstrating a functional role of this pulvinar to V1 communication. These observations are consistent with the hypothesis that the pulvinar modulates the neuronal gain in V1, gating its homeostatic plasticity response.

Given that pulvinar-to-V1 projections can exert an inhibitory effect (*30*), the reduced pulvinar connectivity is consistent with the evidence that monocular deprivation reduces resting GABA in V1(*31*), and that monocular deprivation enhances visual evoked VEP and fMRI responses in the visual cortex (*6, 32*), suggesting that plasticity may be associated with increased cortical excitability. There is also evidence for increased excitability in the visual cortex following short-term binocular deprivation (blindfolding (*33, 34*)), but whether this other form of short-term plasticity is accompanied by decreased pulvinar-to-V1 connectivity is currently unknown. Thus, available evidence is suggestive of a relation between short-term deprivation, shifted excitation/inhibition balance and pulvinar connectivity, but demonstrating the link among these phenomena remains a goal for future research.

We hypothesize that a down-weighting of pulvinar inputs to V1 may be a common trait of diverse adult V1 plasticity phenomena, including monocular deprivation and perceptual learning, as both are associated with reduced pulvinar-V1 connectivity and decreased cortical GABA (*20, 31*). While perceptual learning is accompanied by structural plasticity (*20*) (like motor learning (*35*)), we found that monocular deprivation does not affect structural connectivity (assessed through DWI). This suggests that structural plasticity may require a longer period of altered experience than the mere two hours of our deprivation.

Although the modulation of pulvinar functional connectivity was mainly localized within the visual cortex, sparser but significant connectivity reductions were also observed in auditory and somatosensory areas. This is in line with evidence that monocular deprivation impacts multisensory processing (*36, 37*). Such modulation could be part of a recalibration response, which may be required to preserve the alignment across modalities whenever processing in one sensory modality is altered. It is important to acknowledge that our results do not offer insights into the mechanisms implementing these connectivity changes. Strong functional connectivity can either be supported by direct communication between two regions, or it can be mediated by a third region that is strongly connected with both. This implies that auditory and somatosensory areas may show a deprivation effect due to their common input from cortical, thalamic or brainstem circuitry, leaving open the interpretation of this result.

Based on the knowledge that the pulvinar is crucial for conveying modulatory inputs to early sensory areas (*15, 16, 18*), we suggest that a primary effect of monocular deprivation is to reduce the impact of top-down signals on early visual processing. This is in line with previous evidence of impaired multisensory integration (*36*) and reduced feedback signaling (alpha rhythms) (*32, 37*) after monocular deprivation. The reduction of top-down influences could result from the sensory conflict generated by deprivation, as the occluded eye fails to signal events carried by the other senses. A similar effect would be achieved by distorting the monocular image (*38*), applying a monocular inverting prism (*9*), or monocular delay (*10*), all of which introduce a sensory conflict and mimic the effects of monocular deprivation, despite leaving the strength of the visual stimulation unaltered. Repeated incongruence reduces the integration weights assigned to multisensory signals, thereby allowing for sensory recalibration (*39*). On this view, ocular dominance plasticity in adults may be re-interpreted as a recalibration response, triggered by the reduced top-down modulatory inputs conveyed by the pulvinar to V1. This perspective is in line with suggestions that the pulvinar plays a key role in cortical plasticity and the reshaping of functional organization in the visual cortex (*40*).

To conclude, our results stimulate a new understanding of short-term monocular deprivation effects, which goes beyond local gain-changes of evoked V1 responses, pointing to a functional reorganization of the visual processing network. We speculate that experience-dependent changes in early visual cortex depend on reducing the multimodal influences conveyed by the pulvinar, implying that the pulvinar may be the gatekeeper of adult sensory plasticity.

## Materials and Methods

### Experimental Design

Experimental procedures are in line with the principles of the declaration of Helsinki and were approved by the regional ethics committee [Comitato Etico Pediatrico Regionale—Azienda Ospedaliero-Universitaria Meyer—Firenze (FI)] and by the Italian Ministry of Health, under the protocol ‘Plasticità e multimodalità delle prime aree visive: studio in risonanza magnetica a campo ultra alto (7T)’ version #1 dated 11/11/2015. Written informed consent was obtained from each participant, which included consent to process and preserve the data and publish them in anonymous form. We analyzed data from two fMRI sessions, acquired before and after two hours of monocular deprivation. Twenty-five healthy volunteers with normal or corrected-to-normal visual acuity were tested. Sample size was set based on the minimum number of participants (N = 18) required to reliably detect a medium sized effect of monocular deprivation on fMRI signals(*6, 7*). Due to image distortions, data from 3 and 6 participants were discarded from the first (TR = 3s) and second (TR = 1s) fMRI dataset. This left respectively 22 participants for the TR = 3s fMRI dataset (12 females mean age ± S.E.M.= 26.5 ± 0.82, also included in the analyses of the DWI dataset) and 19 participants for the TR = 1s fMRI dataset (12 females, mean age ± S.E.M.= 26.5 ± 0.84).

### Short-term monocular deprivation

Monocular deprivation was achieved by patching the dominant eye for two hours. Like in previous studies (*5, 6*), we used a translucent eye-patch made of plastic material allowing light to reach the retina (luminance attenuation 0.07 logUnits). During the monocular deprivation, participants were free to walk, read and use a computer, but they did not eat nor sleep.

### Binocular rivalry

Ocular dominance changes (before vs. after deprivation) were evaluated with two brief binocular rivalry sessions, each comprising two 3-minute runs, immediately prior to each fMRI session. The visual stimuli were presented on a 15-inch LCD monitor viewed from 57 cm through anaglyph goggles (blue filter on the right lens, red filter on the left). The stimuli consisted of two oblique gratings (red and blue), oriented at ±45°, measuring 3° in diameter, with a spatial frequency of 2 cycles per degree (cpd) and 50% contrast. Equiluminance between the red and the blue gratings was achieved by reducing the intensity of the red channel based on photometric measurements (mean luminance of both the red and blue grating: 1.8 cd/m2). The dichoptic gratings were surrounded by a binocular sharp-edged white frame (encouraging fusion) and displayed against a uniform black background. During the experiment, participants continuously reported their perception by keeping one of three keys pressed: right or left arrows to report dominance of the clockwise or counter-clockwise oblique gratings; down-arrow key to report mixed percepts. Dominance phase durations shorter than 0.3 seconds (finger errors) and longer than 30 seconds (failed rivalry) were discarded. The dominant eye was defined as the eye showing the longer mean phase duration during a baseline binocular rivalry measurement performed in a separate training session.

We defined ocular dominance as the total time during which the stimulus presented in the dominant eye was perceived, divided by the total time during which the stimulus in either eye was perceived (which amounts to the total testing time minus the time during which a mixture of the two stimuli was perceived):

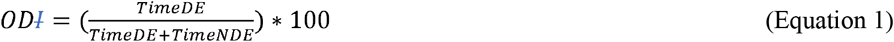

where OD stands for Ocular Dominance, DE for dominant eye and NDE for non-dominant eye. The effect of monocular deprivation was quantified as the ODI difference before and after the two-hour deprivation.

### MR system and sequences

Scanning was performed on a 7T-MRI system (SIGNA7T-PISA, GE Healthcare, Milwaukee, WI, USA) equipped with a 2-channel transmit and a 32-channel receiver head-coil (Nova Medical, Wilmington, MA, USA).

Anatomical images were acquired at 0.8 mm isotropic resolution using a MP-RAGE (Magnetization Prepared RApid Gradient Echo) sequence with the following parameters: echo time TE = 3.5 ms, GRE repetition time TR_GRE_ = 7.3 ms, repetition time TR_MPRAGE_ = 3380 ms, inversion time TI = 1100 ms, delay time TD = 1600 ms, flip angle FA = 8°, rBW = 62.5 kHz, field of view FOV = 220 mm, matrix = 276×276, slice thickness = 0.8mm, 200 slices.

Two sets of functional MR images were acquired at the same isotropic spatial resolution of 1.5 mm (matrix = 128×128, slice thickness = 1.4 mm, slice spacing = 0.1 mm), rBW = 250 kHz, FA=77°, ASSET = 2, phase encoding direction Anterior-to-Posterior (AP).

The first fMRI dataset had TR = 3000 ms, TE = 23 ms, number of volumes = 160.

The second fMRI dataset had TR = 1000 ms, TE = 19.3 ms, number of volumes = 300.

Each functional sequence included 4 initial dummy volumes, to allow the stationarity of the signal. For each EPI sequence, we acquired four additional volumes with the reversed phase encoding direction (Posterior-to-Anterior, PA), used for distortion correction. We acquired each set of functional images (TR = 3s first, followed by TR = 1s) in resting-state with eyes closed.

For the study of structural connectivity, Diffusion Weighted Imaging (DWI) datasets were acquired by using a MUltiplexed Sensitivity-Encoding (MUSE) sequence with 2 mm isotropic resolution (FOV = 200 mm, matrix =100×100, slice thickness = 2 mm). The other acquisition parameters were: TR = 5500 ms, TE = 50.5 ms, 2 echoes, ASSET = 2, 32 noncollinear diffusion gradient directions with b-value of 1000 s/mm^2^, four volumes without b-value = 0 s/mm^2^.

### MRI preprocessing

Preprocessing of anatomical and functional MR images was performed with BrainVoyager (*41*) (version 20.6), Freesurfer (*42*) (v6.0.0), FSL (*43*) (version 6.0.4) and ANTs (*44*). In addition, Matlab (The MathWorks Inc.) was used to perform resting-state analyses.

Anatomical images were processed by a standard procedure for segmentation implemented in Freesurfer (recon-all (*42*)). Transformation matrices from individual participants’ T1 anatomical images to the MNI template (*45*) were computed through an affine registration and a warping procedure using the antsRegistrationSyN.sh routine (*44*).

fMR images were corrected for slice time and motion (*41*) and undistorted using EPI images with reversed phase encoding direction, using the Brain Voyager COPE plug-in (*46*). We then exported the preprocessed images from BrainVoyager to NiFTi format. Each participants’ first functional acquisition (acquired right after the anatomical images) was aligned to the corresponding anatomy using a boundary based registration algorithm (Freesurfer bbergister function). All functional volumes were aligned to the same first functional acquisition with a rigid transformation (using antsRegistrationSyN.sh and antsApplyTransforms.sh), then registered to the MNI space using the affine transformation and warp fields obtained from the anatomical registration (described above, matrices applied with antsApplyTransforms.sh).

fMR images with TR = 3s were preprocessed as follows. After slice-time and motion correction, the fMRI time-series were band-pass temporal filtered (0.01– 0.1 Hz, minimum-order filter with a stopband attenuation of 60 dB); to minimize filtering artifacts, we excluded the first and last 5 TRs from further analysis. Next, we regressed out three main potential confounds (modelled by 14 regressors), as in previous work on pulvino-cortical interactions (*13*): (1) the six motion-parameter estimates and their temporal derivatives, (2) the signal from the three ventricular spaces (average signal in the first, second and third ventricles, defined by the recon-all routine (*42*)) and (3) the signal from the cerebral white matter (defined by the Harvard-Oxford subcortical atlas (*47*)).

fMR images with TR = 1s were preprocessed in a similar way. After slice-time and motion correction, the fMRI time-series were high-pass temporal filtered (0.0078 Hz, minimum-order filter with a stopband attenuation of 60 dB); to minimize filtering artifacts, we excluded the first and last 5 TRs from further analysis. Next, we regressed out three main potential confounds (modelled by 20 regressors): (1) the physiological parameters (respiration and cardiac pulsation preprocessed with the PhysIO Toolbox (*48*)), (2) the signal from the three ventricular spaces (average signal in the first, second and third ventricles, defined by the recon-all routine (*42*)) and (3) the signal from the cerebral white matter (defined by the Harvard-Oxford subcortical atlas(*47*)).

Diffusion weighted images (DWI) were preprocessed with QSIprep (*49*). Preprocessing involved distortion correction, denoising using MP-PCA, motion correction and registration to anatomical template. Tensor fitting and reconstruction of the whole brain tractogram was performed with pyAFQ 1.3.6. The tractogram was generated with n_seeds = 8 using a CSD model and sampling was done from ROI masks. Optic radiations and pulvinar-V1 radiations were segmented based on LGN, pulvinar and V1 masks imported in pyAFQ. After successfully labelling tracts belonging to the optic radiation, outlier tracts were removed as a part of pyAFQ cleaning pipeline. Fractional Anisotropy (FA) profiles for clean bundles were generated after resampling the tract length at 100 points. For each participant we estimated the FA profiles before and after monocular deprivation and compared them with paired t-tests. The associated two-tailed p-values were evaluated against an FDR (*50*) corrected 0.05 threshold.

### Statistical analysis

#### Functional connectivity analysis

Functional connectivity was estimated with both a within-participant and an aggregated subject approach. For the within-participant approach, we computed the Pearson’s correlation coefficient between the fMRI time-series in pairs of ROIs (average time-series of all voxels within the ROI). The correlation coefficients were Fisher-transformed to ensure normality; the resulting values were compared before vs. after monocular deprivation using paired t-tests (two-sided p-values). To check for possible changes in signal strength across time-points, we also computed the Fourier spectrum of the fMRI time-series in each ROI (in the 0.01-0.08 Hz frequency band) and compared its root mean square before and after deprivation using paired t-tests (two-sided p-values).With the aggregate-subject approach, we generated functional connectivity maps over the whole cortical surface. For this, we concatenated the z-scored individual participant’s fMRI time-series and computed the Pearson’s correlation between the average time-series in each ROI and the time-series of each cortical voxels. We also computed voxel-wise differences of Pearson’s correlation coefficients before and after deprivation. We mapped the three coefficients (correlations before and after deprivation, and correlation differences across deprivation) on a 3D rendering of the cortical surface (fsaverage) and applied a spatial smoothing (FWHM = 3 mm). The two hemispheres were then registered to the same symmetric surface (fsaverage_sym) and the two correlation maps averaged. The individual vertex correlation coefficients were associated with two-sided p-values via Studentization (for the pre and post-deprivation values) or using Fisher transformation (for the post-pre differences). The resulting surface maps (shown in Figure 1 and Extended Data Figure 1 and 2) were thresholded after FDR (*50*) correction of the two-sided p-values (FDR-corrected thresholds for the pulvinar ROI: p_pre_ = 2.90 × 10^−3^; p_post_= 2.80 × 10^−3^; p_difference_ = 1.45 × 10^−4^; LGN: p_pre_ = 1.80 × 10^−3^; p_post_= 1.40 × 10^−3^; p_difference_= NaN, meaning that no voxel passed the threshold; V1: p_pre_ = 3.00 × 10^−3^; p_post_= 2.6 × 10^−3^; p_difference_ = 3.68 × 10^−5^) and cluster corrected (cluster size = 50 vertices). We accompanied these maps with outlines of the main clusters displaying robust connectivity at baseline (|r| > 0.2 and cluster = 500 vertices). We also visualized volume maps (Figure 2), where correlations values before and after deprivation were thresholded after FDR(*50*) correction of the p-values (thresholds: p_pre_ = 9.87 × 10^−6^, p_post_ = 9.87 × 10^−6^; no cluster correction).

### Effective connectivity analysis

From the high-temporal resolution fMR images (TR = 1s, average time-series of all voxels within the pulvinar, LGN and V1 ROIs), we estimated effective connectivity with a within-subjects approach using dynamic causal modeling (*26*) in the rDCM toolbox implementation (*51, 52*) (available as part of the TAPAS software collection (*53*)). The rDCM routine returned a value of “negative free energy”, indicative of the trade-off between accuracy and complexity of the model. We compared these values across two models: one representing the fully connected network, i.e. bidirectional connections between pulvinar and V1, LGN and V1 and between pulvinar and LGN, and one without the connections between pulvinar and LGN (which are not supported by anatomical evidence). Performance of the latter was systematically better across all participants; we therefore reported its connectivity estimates (the “A-matrix”) in the main text (Figure 2); Supplementary Figure S4A also reports the (very similar) connectivity estimates for the fully connected model.

The significance of the individual connections was assessed with one-sample t-tests across participants and the associated two-tailed p-values were evaluated against an FDR (*50*) corrected 0.05 threshold.

We further tested the effect of monocular deprivation on effective connectivity with a two-by-two ANOVA for repeated measures, with main factors time (before and after deprivation) and directionality (e.g., pulvinar-to-V1 and V1-to-pulvinar). This was followed up with post-hoc t-tests, and the associated two-tailed p-values were evaluated against an FDR (*50*) corrected 0.05 threshold.

We complemented this DCM analysis with a simpler cross-correlation of the same fMRI time-series. This was implemented with the standard xcorr Matlab function set to the “normalize” option, which returned cross-correlation coefficients normalized by the autocorrelations. To test whether the asymmetry of the curves changed after deprivation, we took the average cross-correlation at the three negative and positive lags closest to zero ([−3 −2 −1] and [+1 +2 +3]) and evaluated their difference before vs. after deprivation with a series of paired t-tests across participants. The associated two-tailed p-values were evaluated against an FDR (*50*) corrected 0.05 threshold.

### ROI definition

All ROIs were defined in the MNI space, to which our functional scans were registered. For each subject and fMRI acquisition, we extracted the average BOLD signal from all voxels within the ROI (using the fslmeants function).

The pulvinar ROI was taken from a published atlas (*21*) defined from Diffusion-Weighted Images.

The Lateral Geniculate Nucleus (LGN) was taken from the NSD dataset, which outlined it based on a combination of functional data (retinotopic mapping experiments) constrained with anatomical features (*25*).

The V1 ROI was taken from the Glasser cortical parcellation atlas (*24*), defined in the fsaverage surface space, and projected to the MNI volume (https://identifiers.org/neurovault.collection:1549).

### Pulvinar sub-ROI definition with visually evoked activations

The same participants took part in an additional experiment to identify visually responsive regions within the thalamus. Visual images were presented with MR-compatible goggle set (VisuaStimDigital, Resonance Technologies, Los Angeles, USA), connected to a computer placed in the control room. The goggles covered a visual field of approximately 32 x 24 deg, with a resolution of 800 x 600 pixels, mean luminance 25 cd/m^2^, frame rate 60Hz. We used similar stimuli as in (*7*): bandpass noise stimuli (peak 0.4 cpd, bandwidth 1.25 octave), refreshing at 8 Hz, presented in a block design with 9 second stimulation followed by 12 seconds of blank screen (except for a 0.5-degree red fixation spot that was always visible). The contrast of the stimuli was variable (6, 12, 25, 50 and 100%); each run included 10 blocks with the five contrasts levels presented in ascending order and repeated twice.

The experiment consisted of a total of four runs with 70 TRs: two runs in passive viewing followed by two runs with attention allocated to the left or right half of the visual field (one side per run) to detect small color-changes (blue blobs with 1 degree radius, presented for four monitor frames, i.e. 67 ms) superimposed to the bandpass noise stimulus and occurring at random times and locations during the 9 seconds of stimulus presentation. Participants counted the number of transients occurring in each block on the cued side, which varied randomly between 2 and 6, and they reported by keypress whether it was smaller or larger than 4.

fMRI time-courses were slice-time and motion corrected. For each voxel within the pulvinar ROI, we computed the BOLD signal variations (in % signal change) with each stimulus presentation, forming epochs of 7 TRs (21 seconds) referenced to the 1st TR. The mean over time in each epoch defined the event-related response. We averaged these event-related responses across the 20 epochs (10 per run) and our 22 participants to evaluate the strength of the BOLD modulation in each voxel of the pulvinar region.

These maps were used to identify maximally responsive voxels in each condition: passive viewing vs. with attention allocated to the stimulus area. We thresholded maps at 0.25% BOLD signal change and defined binary masks of activated voxels after applying a small spatial smoothing (1 voxel full width at half maximum). Voxels responding in passive viewing or both conditions were assigned to a first region, which was ventrally located; voxels selectively responding to attended stimuli were assigned to a second region, which was localized in a more dorsal territory. This produced two non-overlapping masks of 1036 and 1111 voxels (relatively dorsal and ventral regions respectively, with the number representing voxels in both hemispheres), collectively covering 55.4% of the anatomically defined pulvinar ROI (*21*) (3876 voxels across both hemispheres).

## Acknowledgments

The authors would like to thank Paolo Bosco for help with data preprocessing, and David C. Burr for commenting on the manuscript.

## Funding

European Research Council (ERC) under the Horizon EU framework, Consolidator grant PREDACTIVE (n. 101170249)

European Union - Next Generation EU, in the context of The National Recovery and Resilience Plan, Investment 1.5 Ecosystems of Innovation, Project Tuscany Health Ecosystem (THE, CUP I53C22000780001)

European Union - Next Generation EU, grant PRIN 2022 (Project ‘RIGHTSTRESS—Tuning arousal for optimal perception’, Grant no. 2022CCPJ3J, CUP I53D23003960006).

The Italian Ministry of University and Research under the program FARE-2 (grant SMILY).

