## Supplemental figures for "The pulvinar regulates plasticity in human visual cortex"

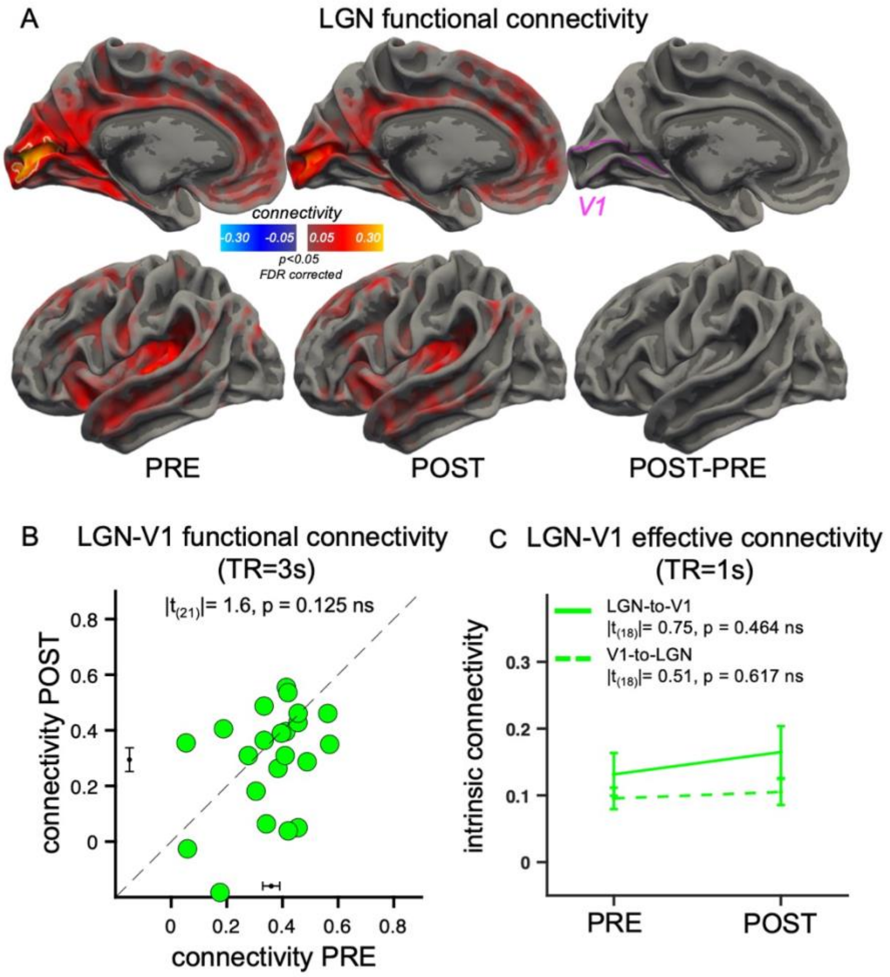


**Fig. S1. LGN functional and effective connectivity**

***A***: *Unaltered functional connectivity of LGN after monocular deprivation (same conventions as in Figure 1A of the main text). In the pre- and post-deprivation maps, connectivity is more focused in the V1 region compared to the pulvinar connectivity (compare with Figure 1A). In the map of connectivity changes, no vertex passes the significance threshold. The white outline in the pre-deprivation map identifies the clusters of regions where connectivity is highly reliable (|r| > 0.20 and 500 vertices), with little spillover outside V1 consistent with anatomy and probably reflecting interareal correlations.*

***B***: *Unaltered functional connectivity between LGN and V1 (atlas-based definition, pink outline) for the individual participants. Pearson's r values were Fisher transformed to allow for statistical comparisons; the black error bars near the axes show the mean +/- S.E.M. connectivity, similar before (0.36 ± 0.03) and after deprivation (0.29 ± 0.04), Cohen's d of the difference = 0.34. The text inset reports the results of the paired t-test comparing functional connectivity before and after* *deprivation (p > 0.05 FDR corrected). A and B show data from the full-volume fMRI acquisition, with TR = 3s.*

***C**: Unaltered effective connectivity (intrinsic connectivity, i.e. A-matrix in rDCM) between LGN and V1 (TR = 1s), shown as mean and S.E.M across participants. Text insets report the post-hoc t-tests comparing effective connectivity values before and after deprivation for the two directionalities.*

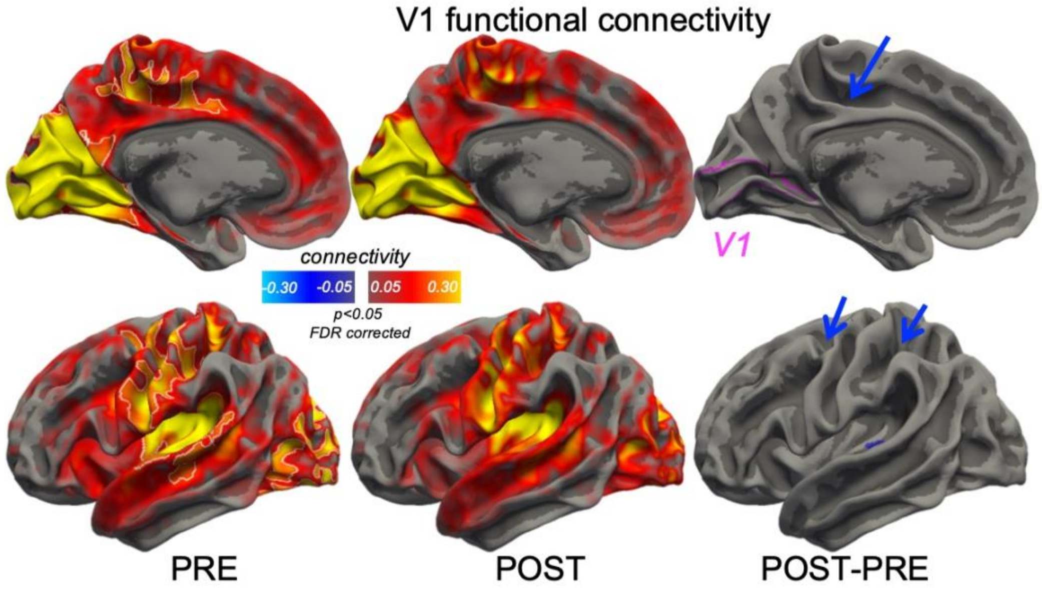


**Fig. S2. V1 cortical functional connectivity**

*Functional connectivity of V1 (atlas-based definition (24), pink outline in the rightmost panels) with the rest of the cortex (same conventions as in Figure 1A of the main text). Connectivity is strongest within the occipital cortex as well as in the auditory and somatosensory networks, similar to the pulvinar connectivity (compare with Figure 1A). After monocular deprivation, there is a selective reduction of connectivity with high-level multimodal areas: cingulate gyrus, superior temporal gyrus, intra-parietal sulcus and area 6a (the four blue spots visible in the rightmost maps). Connectivity is not reduced in occipital regions, and this is consistent with the finding that the post deprivation connectivity of the pulvinar is homogenously reduced in all occipital cortex (implying that the cortical-cortical connectivity in the occipital pole does not change with deprivation). In other words, the only connectivity reduction is the one mediated by the pulvinar: disconnecting the pulvinar from the occipital areas does not change interareal connections.*

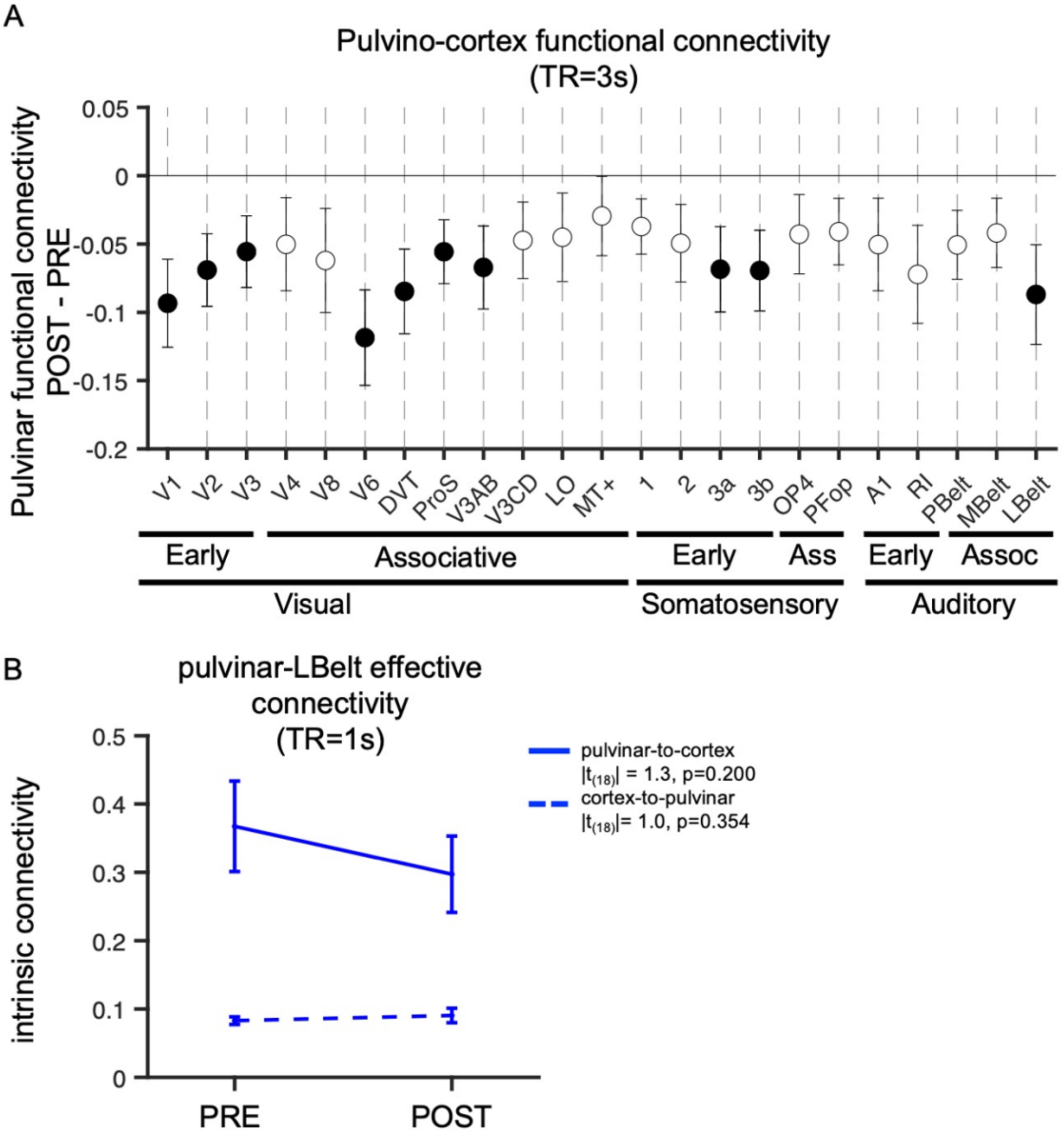


**Fig. S3: Pulvinar connectivity changes beyond V1**

***A.*** *Pulvinar functional connectivity changes (mean and s.e. across participants) measured at TR=3s for areas defined according to the Glasser cortical parcellation atlas (24). Filled symbols are significant changes at $p < 0.05$ uncorrected. Areas V3AB and 3a-b encompass the two areas identified by blue arrows in Figure 1A.*

***B.*** *We followed up this analysis for the LBelt area in the temporal cortex that was also covered in our partial-volume acquisitions at TR = 1 second. A DCM model of the connection between this region and the pulvinar revealed a non-significant trend for reduced effective connectivity in the pulvinar-to-cortex direction, and no trend for reduced cortex-to-pulvinar connectivity (statistics in the text insets). This adds to the evidence in Figure 2 suggesting that communication from the* *cortex to the pulvinar is not affected by monocular deprivation; although the pulvinar keeps receiving its normal cortical input, its projections back to the cortex (especially the visual cortex) become less effective.*

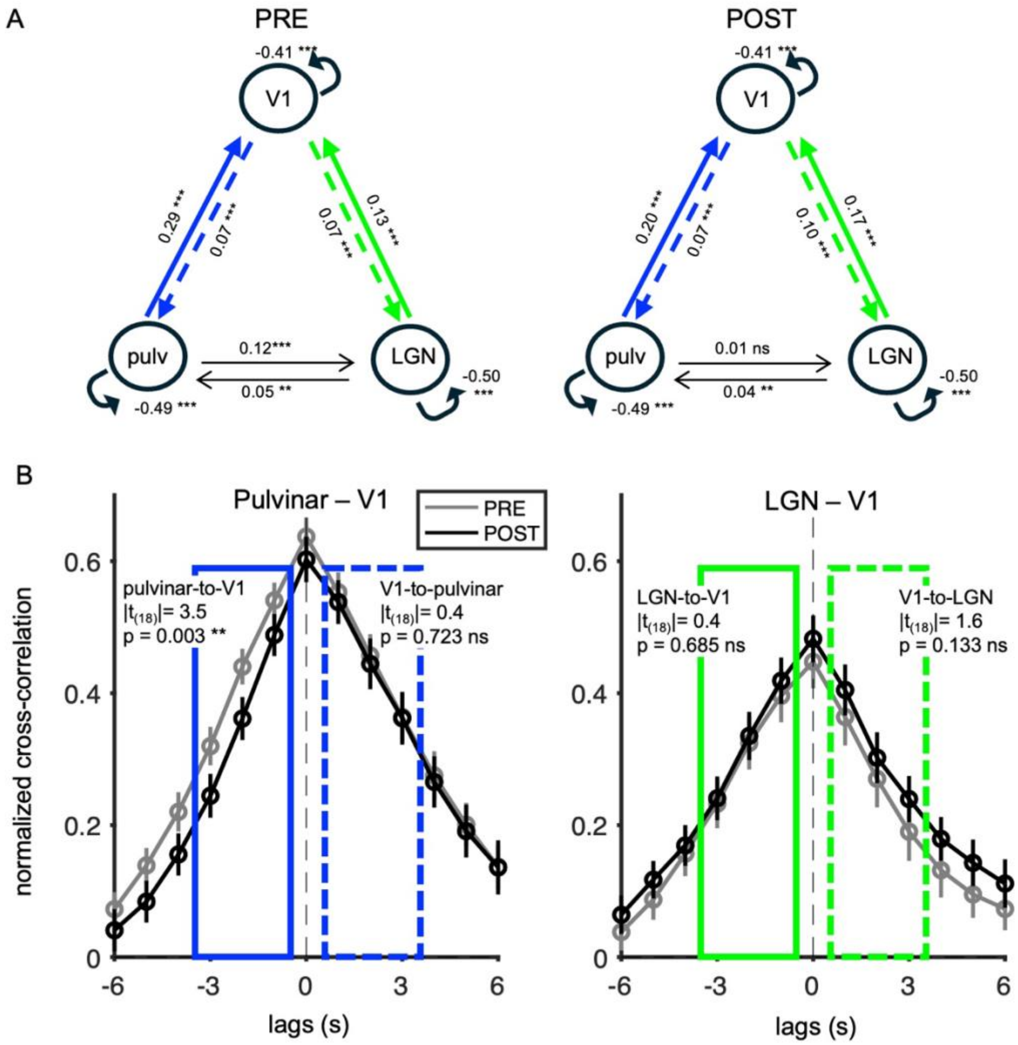


**Fig. S4. Fully connected DCM network and thalamo-cortical cross correlation profiles.**

***A.*** *Alternative network definition for the DCM, assuming full connectivity among all regions. Connectivity estimates between V1 and pulvinar, and between V1 and LGN are remarkably similar as in the simpler model shown in Figure 2 (selected due to its better fit to the data, according to the negative free energy parameter). The fully connected model assigned significant values to the pulvinar-to-LGN connectivity before (not after) deprivation, and a significant ($|t_{(18)}| = 2.62$, $p = 0.017$) decrease of connectivity after deprivation. This connectivity, not validated on anatomical evidence, may reflect a common input to LGN and the inferior pulvinar (its retino-recipient portion, (40)). However, interpretation of this connectivity should be cautious given that this* *connection is not present in the best fitting model (Figure 2). The statistics of the effects of monocular deprivation on each pair of connections are as follows.*

*For connectivity between the pulvinar and V1, the 2x2 ANOVA with factors time (pre vs. post deprivation) and directionality (pulvinar-to-V1 vs. V1-to-pulvinar) showed a significant interaction ($F_{(1,18)} = 12.2$, $p = 0.002$, $\eta^2_{partial} = 0.35$), with a significant decrease of pulvinar-to-V1 connectivity ($|t_{(18)}| = 3.01$, $p = 0.007$, Cohen's $d = 0.69$), and unaffected V1-to-pulvinar connectivity ($|t_{(18)}| = 0.09$, $p = 0.929$, Cohen's $d = 0.02$).*

*For connectivity between the LGN and V1, the 2x2 ANOVA showed no significant interaction ($F_{(1,18)} = 0.6$, $p = 0.457$) and post-hoc t-tests revealed no significant change of either LGN-to-V1 ($|t_{(18)}| = 0.80$, $p = 0.464$, Cohen's $d = 0.17$) or V1-to-LGN ($|t_{(18)}| = 1.66$, $p = 0.114$, Cohen's $d = 0.38$).*

*For connectivity between the pulvinar and LGN, the 2x2 ANOVA showed a significant interaction ($F_{(1,18)} = 4.9$, $p = 0.039$, $\eta^2_{partial} = 0.22$), with a significant decrease of pulvinar-to-LGN connectivity ($|t_{(18)}| = 2.62$, $p = 0.017$, Cohen's $d = 0.60$), and unaffected LGN-to-pulvinar connectivity ($|t_{(18)}| = 0.69$, $p = 0.500$, Cohen's $d = 0.16$).*

***B.*** *Normalized cross-correlation of the average time-series (TR = 1s) for the pulvinar or LGN and V1 ROIs, measured before (gray symbols) and after monocular deprivation (black symbols) and plotted as a function of time-lag. Symbols show mean and S.E.M. across subjects. For the pulvinar-V1 cross-correlation (left), there is a clear reduction at negative lags (estimating the connectivity in the pulvinar-to-V1 directionality) and no effect at positive lags, estimating the connectivity in the V1-to-pulvinar directionality. That values are reduced several seconds before the peak likely resulting from the low-pass behavior imposed by the hemodynamic response function. The significance of these effects was estimated by averaging cross-correlation values at the three time-lags preceding or following 0 (blue boxes with continuous or dashed lines respectively) and comparing them before vs. after monocular deprivation (text insets in each box; the corresponding Cohen's d values are 0.67 and 0.12 for the pulvinar-to-V1 and V1-to-pulvinar directionality respectively). For the LGN-V1 cross-correlation (right, same format but with green colors), no significant changes were observed, neither at negative nor at positive lags (Cohen's d = 0.15 and 0.41 respectively).*

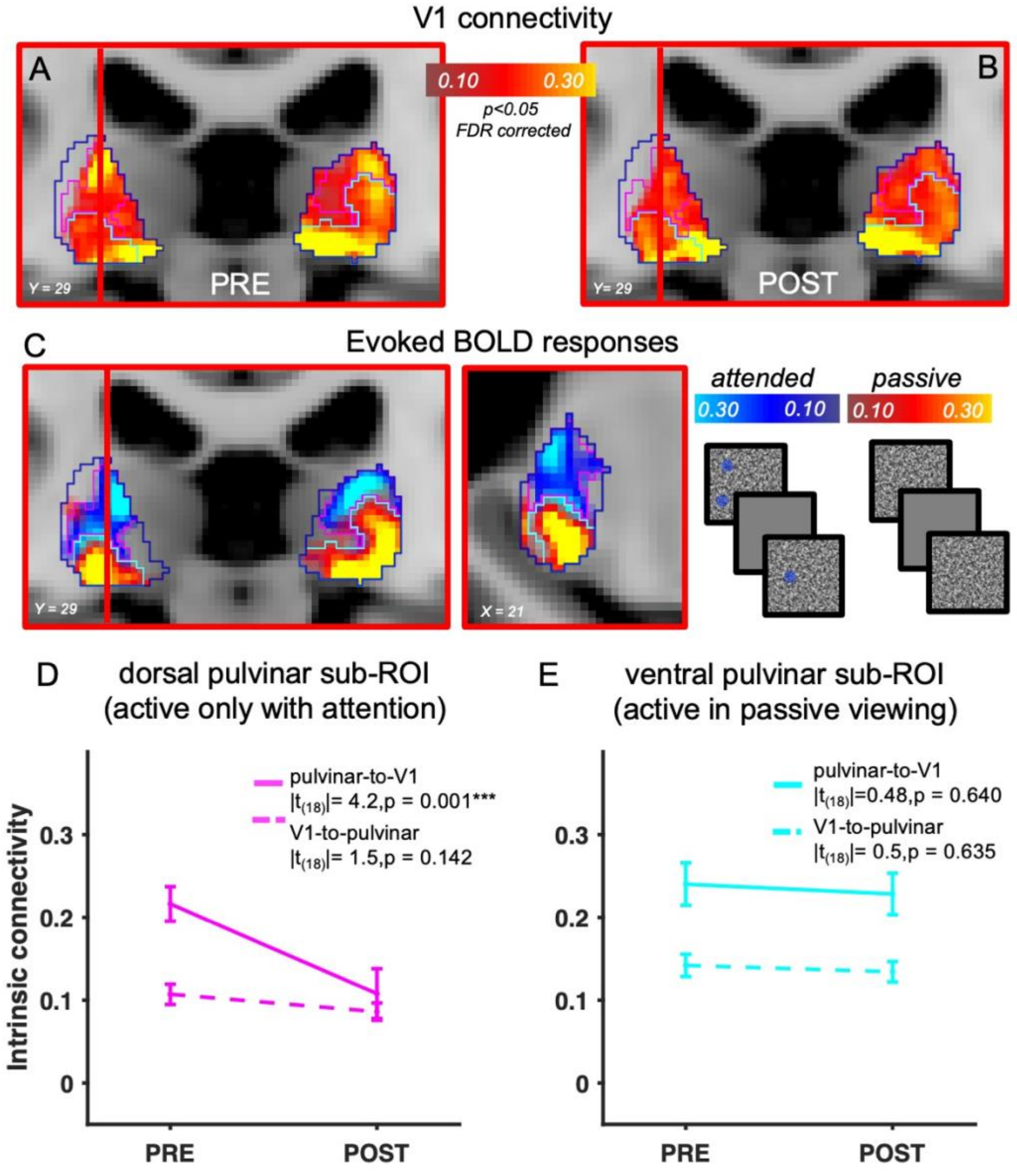


**Fig. S5. Parcellation of the pulvinar based on visually evoked responses.**

***A-B.*** *V1 functional connectivity in the pulvinar ROI, before (A) and after deprivation (B). The magenta and cyan outlines draw the sub-regions of interest defined from the visually evoked activity described in C. The red line identifies the sagittal slice shown in Figure 2C-D of the main* *text. V1 connectivity clustered in two regions: a larger and stronger ventral hotspot and a smaller/sparser dorsal one.*

***C**. Visual evoked activity within the pulvinar ROI, measured in two conditions: passive viewing (activations shown with warm colors) and with attention allocated to the stimulus area (cold colors). Values are % signal change (average BOLD modulation in each of 20 stimulation epochs and across the 22 participants). The magenta and cyan outlines show the two sub-regions of interest defined on these activation patterns: including voxels that responded in the passive viewing condition (cyan, relatively ventral) or voxels that selectively responded when attention was allocated to the stimulus area (magenta, relatively dorsal). The topography of these activation patterns largely matched the topography of V1 connectivity (A-B). However, the match was not exact, as responses in the passive viewing condition (which were stronger in the ventral pulvinar) in some cases also extended into the dorsal hotspots of V1 connectivity (possibly linked with the random allocation of attention in the passive viewing condition). These pulvinar ROI definitions are largely coherent with the definitions used in previous studies on pulvinar connectivity (13,28,23). Although our anatomical mask (21) does not include a mediodorsal region, characterized by strong inferior parietal connectivity (13,28), it does cover most of the regions labelled ventral and dorsal in previous work, and the latter overlaps with our relatively dorsal region defined from functional activations, not with our relatively ventral region.*

***D-E**: DCM results for the two sub-regions of the pulvinar outlined in A-C (same model used in Figure 2, substituting the anatomically defined pulvinar ROI with either of the two functionally defined pulvinar sub-ROI showed in panels A-C). A 2x2 ANOVA with factors time (pre vs. post deprivation) and directionality (pulvinar-to-V1 vs. V1-to-pulvinar) showed a significant interaction for the dorsal sub-ROI ($F_{(1,18)} = 13.3$, $p = 0.002$, $\eta^2_{partial} = 0.43$), but not for the ventral one ($F_{(1,18)} = 0.8$, $p = 0.385$, $\eta^2_{partial} = 0.04$). Text insets show the results of post-hoc t-tests, with the same format as in Figure 2F. In no case did connectivity between V1 and LGN changed (all $|t|<1.4$, $p > 0.170$).*
